# In vitro evaluation of the decontamination effect of cold argon plasma on selected bacteria frequently encountered in small animal bite injuries

**DOI:** 10.1101/353821

**Authors:** S. Winter, A. Meyer-Lindenberg, G. Wolf, S. Reese, M.C. Nolff

## Abstract

**Objective:** The beneficial effects of cold argon plasma (CAP) on wound healing and its capacity for bacterial decontamination has recently been documented. However, despite favourable reports from experimental trials and human applications, the first in vivo studies in small animals did not prove any decontamination effect in canine bite wounds.

The present study therefore aimed to evaluate the decontamination effect of CAP in different bacteria frequently encountered in canine bite wounds *in vitro.*

**Methods:** Standard strains of *Escherichia (E.) coli, Staphylococcus (S.) pseudintermedius, S. aureus, Streptococcus (Sc.) canis, Pseudomonas (P.) aeruginosa* and *Pasteurella multocida* were investigated. To evaluate the influence of the bacterial growth phase, each bacterium was incubated in nutrient broth for 3 and 8 hours, respectively, before argon plasma treatment. Three different bacterial concentrations were created per bacterium and growth phase, and each was exposed to cold plasma at a gas flow rate of 5 standard litres/minute of argon for 30 seconds, 1 minute and 2 minutes.

**Results:** Argon treatment resulted in acceptable decontamination rates (range 98.9-99.9%) in all bacteria species in vitro; however, differences in susceptibility were detected in the different tested bacteria. Treatment time significantly (P<0.05) correlated with the decontamination rate in *E. coli*, *Sc. canis* and *S. aureus*, with an exposure time of 2 minutes being most effective. The initial bacterial concentration significantly (P<0.05) influenced decontamination in *Pasteurella multocida* and *P. aeruginosa,* in which treatment time was not as important. The growth phase only influenced decontamination in *S. pseudintermedius.*

**Conclusion:** CAP exerts effective antibacterial activity against the tested bacteria strains in vitro, with species specific effects of treatment time, growth phase and concentration.

## Introduction

Microbial multidrug-resistance (MDR) is one of the main issues that must be solved by modern medicine [1,2,3]. Among surgical patients, surgical site infections, and especially open wounds, represent risk groups, and antibiotic treatment frequently results in a shift to more resistant bacteria rather than in wound decontamination [1,3,4]. Recognition of this problem has increased research on alternative strategies for wound decontamination, including antiseptic substances and physical treatment options in human medicine [5,6]. However, despite the fact that the same problem exists in small animal surgical patients [1,3], there is a paucity of studies focusing on antiseptic wound treatment in veterinary medicine. Numerous in vitro and in vivo trials in small mammals and humans have documented an effective decontamination effect combined with a wide safety margin regarding a damage of eukaryotic cells for CAP [7]. Three major factors lead to bacterial cell destruction [20, 21]:

1. Destruction of the bacterial cell membrane
2. Intracellular protein damage
3. Direct DNA damage

Loaded particles, such as ions and electrons, create an electrostatic field that permeates the bacterial cell wall [22]. The electrostatic force accumulates on the cell surface until it causes the outward directed cell wall force to subside, leading to destruction [20, 22, 23]. This effect is more profound in gram negative bacteria, due to their more irregular rough outer membrane. While morphologic cell wall destruction has been documented in gram negative bacteria species, no morphological changes occur in gram positive bacteria [22, 23].

In addition, oxidative stress exerts direct effects on proteins and bacterial DNA and results in damage of the cell wall as well as intracellular components [24–27]. Massive oxidative stress results in lipid-oxygenation, loss of cellular cytoplasm, proteins and oxidative damage to the DNA, leading to cell death once the reparation mechanism is overcome [27–31].

Because the effect of plasma is related to numerous factors, a unique ‘dosing’ protocol is not available [24,26]. The type of effluent is closely related to the carrier gas, the room air and the device used for generation of the plasma state [24, 26]. Furthermore, the distance between the plasma source and the wound surface influences the decontamination effect [32]. Longer treatment times have been found to result in greater ‘bacterial death’ however; this is influenced by bacteria species, growth form (biofilm) and substrate on which the bacteria grow [23,32–35]. Factors such as bacterial growth state (growing or steady state) and initial bacterial concentration have not been investigated so far.

Despite these well-investigated facts, the first clinical trials evaluating the decontaminating effect of CAP in canine bite wounds were not able to detect any decontamination effects in vivo [36]. Therefore it was our aim to investigate the effect of the used CAP source in bacteria to be anticipated in canine bite wounds in vitro in order to document the in vitro efficacy and potential influencing factors with respect to bacteria species, bacteria growth phase and bacteria concentration. Based on a recent survey on the bacterial bio-burden of open treated wounds in small animal patients, the following bacteria were chosen for evaluation: *Escherichia (E.) coli, Staphylococcus (S.) aureus, Staphylococcus (S.) pseudintermedius*, *Streptococcus (Sc.) canis*, *Pasteurella multocida* and *Pseudomonas (P.) aeruginosa* [1]

Our hypothesis was that CAP would efficiently decontaminate all chosen isolates with an exposure time of 2 minutes being most effective. Furthermore, we postulated that this effect would be independent from the growth phase of the tested bacterium, and that higher bacterial concentrations would need longer exposure times for decontamination. The aim was to investigate the efficiency of CAP in vitro on selected bacteria before using it in a following in vivo study on canine bite wounds.

## Materials and Methods

### Bacteria selection and culture

The following bacteria were included for further validation: *E. coli* DSM 1103, *S. aureus* DSM 1104, *S. pseudintermedius* DSM 21284, *Sc. canis* DSM 20715, *Pasteurella multocida* DSM 5281 as well as *P. aeruginosa* DSM 1117. All strains were obtained from the Leibniz Institute DSMZ German Collection of Microorganisms and Cell Cultures. To exclude the potential influence of different genetic alterations in wild-type bacteria, standard strains were chosen. Bacterial suspensions were prepared using a standard-I-bouillon (No. 1.07882.0500, Merck) at a pH of 7.5 ± 0.2 at 37°C and plated on Mueller-Hinton Agar (No. 1.05435.0500, Merck). In case of Sc. canis and *Pasteurella* multocida, Mueller-Hinton Agar was supplemented with 5% sheep agar.

### Determination of the bacterial growth phase

Bacterial proliferation can be divided into an exponential growth phase with high metabolic activity followed by a stationary phase. In order to detect differences of the effect of CAP with respect to the growth phase of the investigated bacteria, the transition from growth to stationary phase of each bacterium was determined by visual control using the MC Farland standard. Each enrolled bacterium was incubated in standard I-bouillon at 37°C ± 1°C for 48 hours with a visual control every 2 hours to document the increase of turbidity until the stationary phase was reached (no more detectable increase). Based on the results of the detected change in turbidity, an incubation of 3 hours was representative of exponential growth in all tested bacteria, while an incubation of 8 hours was representative of the stationary phase.

### Determination of initial bacterial concentration

For determination of the colony-forming units per square centimetre (CFU/cm^2^) a log10 serial dilution of the inoculum for plating was used. Colony counting was done after incubation at 37°C ± 1°C for 48 hours. For treatment with Argon plasma, three concentrations (10^-1, 10^-2 and 10^-3) were plated per strain and per growth phase (6 plates per strain) on Mueller-Hinton Agar. After treatment with argon plasma, the plates were incubated at 37°C ± 1°C before counting colonies in the treated areas.

### Cold Argon Plasma Treatment

The plasma device used in this study (KinPen Vet-NeoPlas) was an atmospheric plasma jet with a handheld unit, and consists of a 1 mm pin-tip electrode mounted in the centre of a quartz capillary (1.6 mm inner diameter). Argon was selected as a working gas at a flow of 5 standard litres per minute. The gas is ignited at the tip of the electrode and creates a jet-like effluent covering approximately 1 cm^2^. At these settings, the effluent has a visible length of 13 mm and a constant temperature of 48°C at the tip (Fig 1).

**Fig 1.**
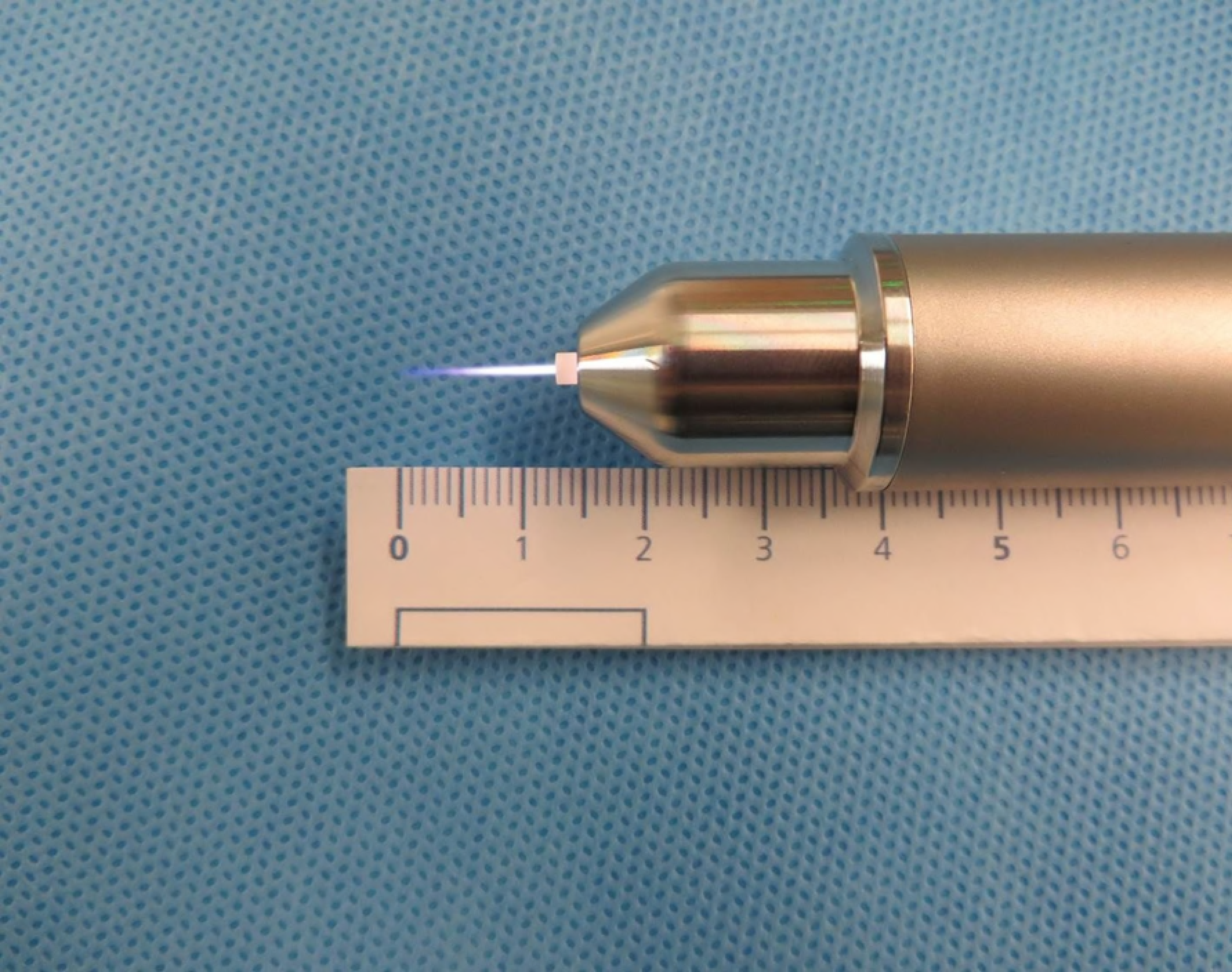
Live view of the KinPen^®^ Vet with ignited plasma beam.

To assure repeatability of treatment, a template was created and placed on the backside of each plate for plasma irradiation (Fig 2). Three areas of 1 cm^2^ each were treated with the plasma device at an approximate distance of 1 cm above the surface for 30 seconds, 1 minute or 2 minutes.

**Fig 2.**
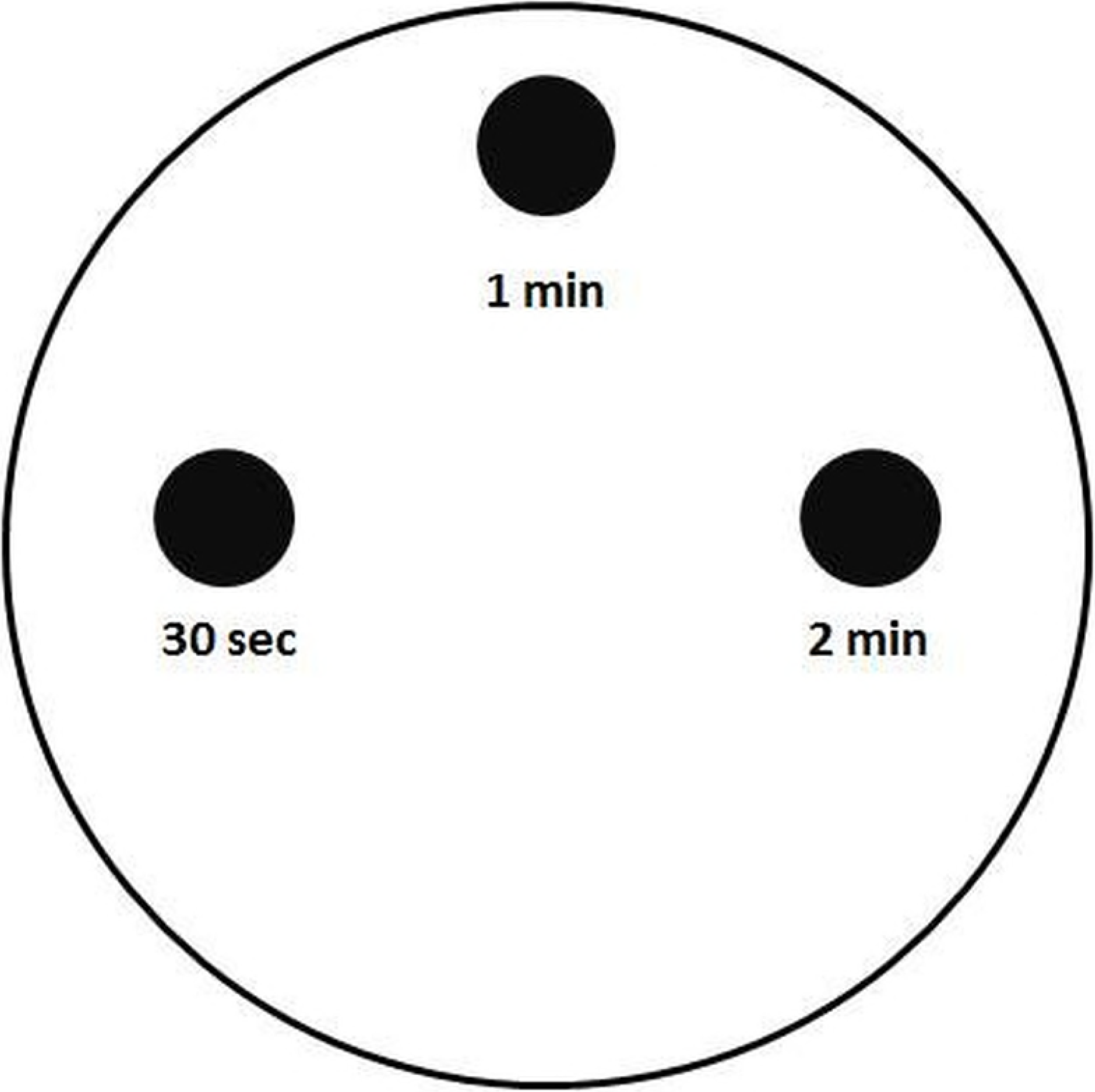
CAP treatment was performed with the KinPen Vet^®^ held in upright position over the well and oriented based on a stencil positioned at the backside of the well.

After the plasma treatment, plates with isolates of the six bacterial strains were incubated for 14 to 20 hours at 37°C ± 1°C. *Sc. canis* was incubated for 44 to 46 hours due to the small size of colonies detected, in order to facilitate easier identification of CFUs. Examination was done twice for each bacterial strain and growth phase. Three different log10 dilutions for each bacterial proliferation were plated respectively on agar plates. Each agar plate had three areas for CAP treatment, achieving 18 areas for single examination of one bacterial strain. Altogether, 36 areas (n=36) for one and 216 areas (n=216) for all six different bacterial strains were examined. This resulted in different concentrations between the two tests of the bacterial strains and also for each growth phase.

### Calculation of the decontamination effect

The CFU per plasma treatment zone of 1 cm^2^ was divided by the CFU per 1 cm^2^ without treatment, calculated from the log10 dilution of the inoculum. The values were given in percentage of growth reduction as well as in log reduction.

### Statistics

Data were computed in Excel (Microsoft Excel 2010) and analysed using SPSS (IBM SPSS Statistics 23.0) software. Descriptive statistics were given as mean and standard deviation. Effect of treatment with respect to the investigated factors was determined using multivariate analysis of variance (ANOVA, SS Type III) and Fisher’s f-test. Treatment time of CAP, bacterial proliferation and initial bacterial concentration were tested regarding their effects on the decontamination. Significance was set at P <0.05

## Results

### Decontamination effect with regard to treatment time

Treatment time represents one of the major aspects of CAP efficiency. The effect of treatment time especially influenced the decontamination of *E. coli*, *S. aureus* and *Sc. canis*. In all three bacterial species, significant effects could be detected (Table 1). Bacterial decontamination was observed in all investigated species at both growth phases. The most efficient tested treatment time was 2 minutes per cm^2^. *p. aeruginosa* and *Pasteurella multocida* were the bacteria least affected by treatment time differences (Table 1).

**Table 1.** Decontamination rate with respect to treatment time.

| Bacteria species | Mean decontamination rate |  |  |  |  |  |  |  |
| --- | --- | --- | --- | --- | --- | --- | --- | --- |
|  | % decontamination |  |  | Log reduction |  |  |  |  |
|  | 0.5 min | 1 min | 2 min | 0.5 min | 1 min | 2 min | P-value | η² |
| *E. coli* | 99.98 | 99.99 | 100 | 4.52 | 5.74 | 5.92 | 0.001 | 0.286 |
| *P. multocida* | 89.97 | 99.98 | 99.99 | 3.97 | 5.15 | 5.16 | 0.096 | 0.084 |
| *P.s aeruginosa* | 98.71 | 99.99 | 100 | 3.64 | 4.37 | 5.59 | 0.227 | 0.045 |
| *Sc. canis* | 83.11 | 97.09 | 99.96 | 2.49 | 3.66 | 3.95 | 0.009 | 0.195 |
| *Staph. aureus* | 72.25 | 79.98 | 97.90 | 1.85 | 2.54 | 4.18 | 0.001 | 0.291 |
| *Staph. pseudintermedius* | 67.85 | 68.19 | 78.31 | 2.29 | 2.40 | 2.99 | 0.413 | 0.021 |

In contrast, the treatment time exerts a major effect on decontamination in *S. aureus*. Treatment duration correlated with the size of the inhibited area, despite the fact that fixed areas were treated in all cases (based on the template). Longer treatment time, especially with 2 minutes per cm^2^, increased the zone of inhibition in all tested bacteria without any need to increase the treatment area (Fig 3).

**Fig 3.**
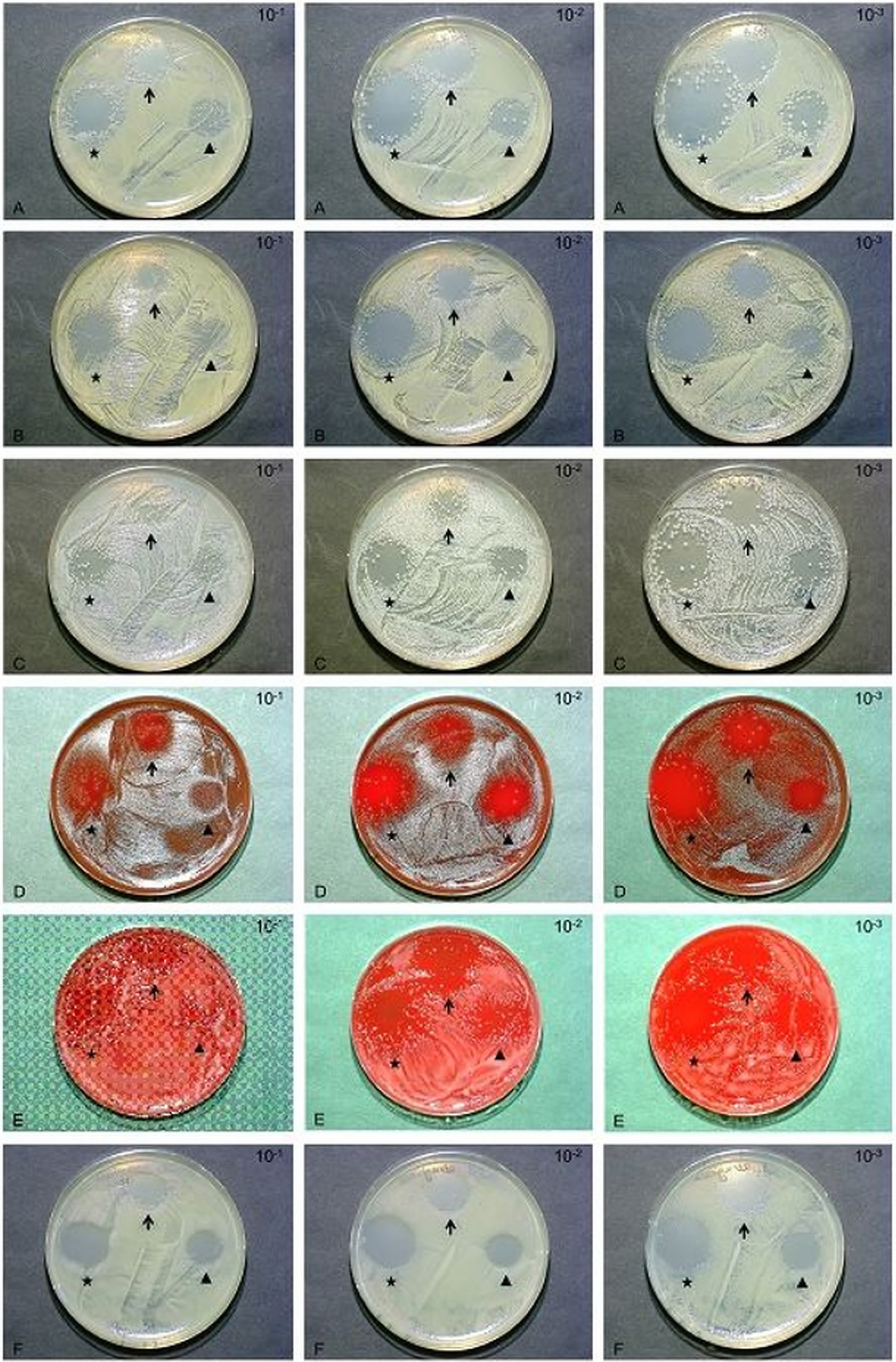
Effect of treatment time (arrowhead 30 seconds, black arrow 1 minute, * 2 minutes) on different concentrations (left row 10^-1, middle row 10^-2 and right row 10^-3 bacteria) of *Escherichia coli* (A), *Staphylococcus aureus* (B), *Staphylococcus pseudintermedius* (C), *Streptococcus canis* (D), *Pasteurella multocida* (E) *as well as Pseudomonas aeruginosa* (F).

### Decontamination effect with regard to bacterial growth phase

The decontamination rate per growth phase and bacterial strain is shown in table 2. Higher decontamination rates were achieved in growing bacteria of *Pasteurella multocida*, *P. aeruginosa*, *S. aureus* and *S. pseudintermedius* as opposed to their stationary forms. However, this difference was only significant for *S. pseudintermedius* (P= 0.001). Only *Sc. canis* showed a slightly higher decontaminating effect in the stationary form. In *E. coli*, no difference was detectable between the two bacterial proliferation phases.

**Table 2:** Decontamination rate with respect to bacterial growth phase.

| Bacteria species | Mean decontamination rate |  |  |  |  |  |
| --- | --- | --- | --- | --- | --- | --- |
|  | % decontamination |  | Log reduction |  |  |  |
|  | growing | stationary | growing | stationary | P-value | η² |
| *E. coli* | 99.99 () | 99.99 | 5.31 | 5.48 | 0.944 | 0.000 |
| *P. multocida* | 97.20 | 96.10 | 4.68 | 4.84 | 0.879 | 0.001 |
| *P.s aeruginosa* | 99.98 | 99.14 | 4.53 | 4.54 | 0.718 | 0.004 |
| *Sc. canis* | 92.49 | 94.28 | 3.09 | 3.64 | 0.373 | 0.025 |
| *Staph. aureus* | 87.66 | 79.09 | 3.18 | 2.54 | 0.428 | 0.020 |
| *Staph. pseudintermedius* | 84.15 | 58.75 | 3.31 | 1.81 | 0.001 | 0.295 |

### Decontamination effect with regard to initial bacteria concentration

The decontamination rate per initial concentration and bacteria species is given in table 3. With the exception of *E. coli*, bacteria concentration significantly influenced decontamination in all tested bacteria. This implies that with higher dilution, more effective decontamination is achieved through lower bacteria concentration. Furthermore the effect of initial concentration seems to be more profound compared with the growth phase of *S. pseudintermedius*. In *Sc. canis*, initial concentration has a more profound effect compared to treatment time.

**Table 3:** Decontamination rate with respect to bacterial concentration.

| Bacteria species | Mean decontamination rate |  |  |  |  |  |
| --- | --- | --- | --- | --- | --- | --- |
|  | % decontamination |  | Log reduction |  |  |  |
|  | growing | stationary | growing | stationary | P-value | η<sup>2</sup> |
| *E. coli* | 99.99 | 99.99 | 5.31 | 5.48 | 0.944 | 0.000 |
| *P. multocida* | 97.20 | 96.10 | 4.68 | 4.84 | 0.879 | 0.001 |
| *P.s aeruginosa* | 99.98 | 99.14 | 4.53 | 4.54 | 0.718 | 0.004 |
| *Sc. canis* | 92.49 | 94.28 | 3.09 | 3.64 | 0.373 | 0.025 |
| *Staph. aureus* | 87.66 | 79.09 | 3.18 | 2.54 | 0.428 | 0.020 |
| *Staph. pseudintermedius* | 84.15 | 58.75 | 3.31 | 1.81 | 0.001 | 0.295 |

### Decontamination effect with regard to bacteria species

The overall rate of bacteria decontamination in all treated areas (n=216) ranged between 71.4 and 99.9%. Treatment of gram-negative bacteria (*E. coli* 99.99% (n=36), *Pasteurella multocida* 96.65% (n=36), *P. aeruginosa* 99.6% (n=36)) resulted in an improved decontamination compared to gram-positive species (*Sc. canis* 93.4% (n=36), *S. aureus* 82.1% (n=36) and *S. pseudintermedius* 71.4% (n=36)).

## Discussion

The aim of this study was to investigate the in vitro efficiency of CAP on selected bacteria frequently encountered in canine bite wounds. Based on the overall effect, CAP treatment seems to represent an interesting alternative to antiseptic wound treatment, especially since no known mechanisms for bacterial resistance have been documented until now (37–39]. Flynn et al. (2015) even documented a high efficiency against so-called ESKAPE bacteria (***E**nterococcus faecium, **S**taphylocoocus aureus, **K**lebsiella pneumoniae, **A**cinetobacter baumannii, **P**seudomonas aeruginosa, **E**nterobacteriaceae*) [17], a group of pathogens frequently encountered in hospital-acquired infections in humans and animals [1,3].

However, the first study evaluating its effect in bite wounds in dogs did not find any decontaminating effect in vivo at all [36]. Based on the beneficial results in experimental and human applications, this finding seems odd. We therefore wanted to evaluate the effect of the plasma source used in vivo in dog bite wounds in an in vitro setting to evaluate the general efficacy. We were able to prove that several factors impact CAP performance, including initial bacterial contamination, treatment duration, bacterial growth phase and bacterial species.

The bacterial contamination of wounds is something that cannot be influenced as it occurs before presentation. Nevertheless, this study underlines the importance to reduce the contamination rate of wounds as far as possible before application of CAP in order to achieve the maximum effect. In a clinical scenario, this can be achieved by debridement and pre lavage, which should always be performed based on our in vitro results.

As previously reported by other authors, we could also detect a superior decontamination effect on gram-negative bacteria compared with gram-positive isolates [20,22,23]. Laroussi et al. (2003) previously found that CAP treatment induced cell wall destruction, leading to loss of integrity and cytoplasmic condensation in gram-negative bacteria compared to untreated controls. In contrast, no differences between treated and untreated gram positive bacteria were detectable [7]. These findings are in concordance with findings documented by Matthes et al. (2013), who reported lower decontamination rates for *S. epidermidis* compared to *P. aeruginosa* [15]. The accumulation of the electrostatic force created by the CAP treatment is more profound in gram-negative species [7,22,].

To our knowledge, there are no other studies in the available literature investigating the influence of the growth phase of CAP treated bacteria. It seems logical, that improved metabolism, as can be encountered in growing germs, might be linked with increased susceptibility to CAP compared to a more resting, static state. We found that this aspect is negligible for most gram-positive and gram-negative tested bacteria species, with the exception of *S. pseudintermedius,* in which growth significantly improved the CAP effect. This finding is interesting, especially as no such difference was detected in *S. aureus*. However, individual differences, even between strains of methicillin-resistant *S. aureus* (MRSA) have been recently described [15].

We also detected an impact of bacteria concentration on decontamination with CAP in all tested bacteria despite *E. coli*, with higher concentrations being less susceptible than lower concentrations. This effect was especially profound in *S. pseudintermedius* and *Sc. canis*. In contrast to bacteria species and growth phase, bacteria concentration can actually be influenced in clinical cases by debridement and intense lavage, which should be included to reduce the concentration and treatment time and increase the efficiency of CAP in clinical situations.

Finally, we found that increased treatment time results in increased decontamination, with the first decontamination effects visible after 30 seconds and a treatment time of 2 minutes being most effective. This is in concordance with the results of Matthes et al. (2013), who described a time-dependent linear efficiency of decontamination between 30 seconds and 5 minutes (300 seconds) [19]. Dose represents a term that remains under debate for plasma applications [24]. Several studies have defined treatment time as dose-; however, the energy level and amount of energy transferred should ideally also be included in determining the effective dose [24]. In addition, adjunctive treatments can increase the effective dose. It has been proven that irradiation of saline induces ionisation in the liquid, lowers the PH and alters reactive species within the fluid, making it bactericidal [24]. Thus, the addition of saline lavage before treatment might exert an even bigger effect than pure CAP treatment.

It is important to consider that CAP dose exerts an impact on bacteria as well as on host cells. Therefore, the maximum dose has to be limited below the threshold that becomes toxic for the host [24]. It has been proven that lower doses of plasma increase angiogenesis in host tissue, while higher doses induce cell death [24,26]. Fortunately, the lethal dose for bacteria is much smaller than the lethal dose for host tissue cells, leaving a wide safety margin for application [9–11,13, 14,16, 18,19, 43].

Based on the profound results achieved in this in vitro trial, we are not able to explain the poor performance of CAP in the clinical situation. We were able to prove, that the used device in the used setting is capable of effectively killing the most frequently encountered bacteria types. However, the data of the current trial underline the importance of appropriate exposure time. The design of the KinPen Vet is adapted to clinical use, facilitating easy application and transportability. Unfortunately, the design allows only treatment of a relative small area. Therefore, a needed dose of 2 minutes per cm^2^ might result in a very long treatment time in clinical patients, prolonging anesthesia time and thus increasing the risk for surgical site infections. We think that this effect represents the most important clinical limitation. If the results of the study by Winter et al. (2018) [36] are interpreted under the light of our current findings, we think that the usage of bigger devices, which allow appropriate treatment time under clinical conditions, would most likely improve performance in vivo.

## Conclusions

CAP exerts a substantial decontamination effect on bacteria frequently found in animal bite injuries in vitro. Because of the lack of bacteria resistance, this treatment might present an interesting alternative for local wound decontamination in vivo. However, bacteria concentration should initially be reduced as far as possible before initiation of CAP treatment in clinical cases. Furthermore, adequate treatment time must be ensured.

## Acknowledgements

The authors wish to thank NeoPlas Greifswald, who supplied the KinPen Vet^®^ used in this study free of charge.

## Conflict of interest

NeoPlas played no role in the study design, data collection, analysis and interpretation of data, or the decision to submit the manuscript for publication. None of the authors has any financial or personal relationships that could inappropriately influence or bias the content of the paper.

